# Structural and mechanical remodeling of the cytoskeleton maintains tensional homeostasis in 3D microtissues under acute dynamic stretch

**DOI:** 10.1101/780312

**Authors:** Matthew Walker, Pauline Rizzuto, Michel Godin, Andrew E. Pelling

## Abstract

When stretched, cells cultured on 2D substrates share a universal softening and fluidization response that arises from poorly understood remodeling of well-conserved cytoskeletal elements. It is known, however, that the structure and distribution of the cytoskeleton is profoundly influenced by the dimensionality of a cell’s environment. Therefore, in this study we aimed to determine whether cells cultured in a 3D matrix share this softening behavior and to link it to cytoskeletal remodeling. To achieve this, we developed a high-throughput approach to measure the dynamic mechanical properties of cells and allow for sub-cellular imaging within physiologically relevant 3D microtissues. We found that fibroblast, smooth muscle and skeletal muscle microtissues strain softened but did not fluidize, and upon loading cessation, they regained their initial mechanical properties. Furthermore, microtissue prestress decreased with the strain amplitude to maintain a constant mean tension. This adaptation under an auxotonic condition resulted in lengthening. A filamentous actin cytoskeleton was required, and responses were mirrored by changes to actin remodeling rates and visual evidence of stretch-induced actin depolymerization. Our new approach for assessing cell mechanics has linked behaviors seen in 2D cultures to a 3D matrix, and connected remodeling of the cytoskeleton to homeostatic mechanical regulation of tissues.

## Introduction

With every breath, heartbeat and movement, cells in our body experience cyclic mechanical stretch, which in turn, creates continually unsteady forces at focal adhesions, across the cell membrane, along cytoskeletal filaments and through the nucleus ^1,2^. In a cell, these forces direct functional and phenotypic behaviors by generating conformational changes, and thereby, alter ligand-receptor affinities^1,2^. Importantly, this ability of cells to feel and adapt to mechanical forces has been linked to crucial events in normal development and function, as well as disease progression, including bone, muscle, heart and lung disorders, and cancer ^3,4^.

In particular, the well-conserved structural elements that make up the cytoskeleton of eukaryotic cells are in themselves mechanosensitive; in response to dynamic stretch, the cytoskeleton softens (decreased elasticity) and becomes more fluid-like ^5–8^. Then upon stretch cessation, it slowly regains its stiffness and resolidifies ^7^. Currently the molecular mechanism(s) behind strain softening remains unclear as there is limited visual-based evidence quantifying cytoskeletal remodeling following cyclic stretching ^9–11^. Furthermore, whereas softening and fluidization has been observed in response to deformation at the subcellular ^5,6^ and single cell levels ^7,8^, the extent of the response remains poorly understood at the tissue-level ^12–14^. Nevertheless, in the body, this response has been linked to the maintenance of airway caliber ^12,13^ and the regulation of blood pressure ^15^, but for unknown reasons, it is absent in certain pathological disorders. For example, unlike in healthy lungs, stretch from a deep inspiration does not dilate asthmatic airways ^16^.

While studying the mechanical behaviors of intact tissues has furthered our knowledge of how cells respond to stretch ^12,13^, these methods possess poor resolution for elucidating subcellular remodeling. Investigations may also be hindered by inter-donor variability and low availability of tissue samples. On the other hand, subcellular cytoskeletal remodeling responses and mechanotransduction pathways have been investigated in large part by growing cells on rigid, flat surfaces. Yet it is known that the physical environment in which a cell is grown alters its mechanical properties and behavior. For example, cells grown on stiff substrates tend to have their actin cytoskeleton arranged into dense stress fibers, and are stiffer, more solid-like and under greater pre-stress when compared to cells on softer substrates ^17–19^.

In addition to matrix stiffness, it is suspected that the mechanical behavior of cells may be further altered by the dimensionality of their environment. In support of this growing hypothesis, culturing cells on a 2D substrate vs. within a more physiologically relevant 3D matrix fundamentally changes the distribution and structure of the cytoskeleton by forcing un-natural apical-basal polarity of adhesion complexes ^20^. The difference between a rigid, flat, petri dish and a soft 3D extracellular matrix (ECM) may also explain observed disparities in cellular behavior, and the loss of efficacy in costly clinical trials that often occurs when pharmaceutical treatments are developed using conventional 2D cell culture techniques ^21–24^. Thus, there exists a need for new high-throughput cell culture techniques capable of probing cell mechanical behavior while maintaining a physiologically relevant soft 3D environment.

To address this need, techniques that allow assessment of the mechanical behavior of cells within reconstituted 3D collagen gels have been a keen interest to the fields of mechanobiology, pharmacology, and tissue engineering ^25^. These methods have furthered our understanding of how tissue-level forces are collectively generated by cells and the ECM ^26–28^. In regards to their response to stretch, it is known that cells within 3D cultures respond to quasi-static changes in matrix tension through altering their contractility in the opposite direction so to maintain tensional homeostasis throughout the cell culture ^29,30^. In other publications these behaviors following step length changes have been linked to actin depolymerization and subsequent reinforcement responses ^31,32^. Although the responses to static strains have been well documented ^29–32^, the mechanical and cytoskeletal remodeling responses under cyclic stretching in 3D cultures are less clear. That said, during dynamic stretching of 3D cultures, the peak force of subsequent loading cycles has been shown to decrease towards a plateau ^33^, which is suggestive of an adaptive strain softening behavior. However, our field lacks a complete characterization of this mechanical response to dynamic stretch in 3D cell cultures and an understanding of how the cytoskeleton structurally remodels.

In addition, the centimeter scale of the bulk gels used in previous investigations limits the experimental throughput, causes imaging difficulties, produces a high diffusive barrier for nutrients and may slow dynamic responses to soluble factors. These limitations of bulk 3D cell cultures can largely be overcome by shrinking the cell culture size through adopting a Lab-on-a-chip approach. For example, Legant *et al.’s* (2009) Microfabricated Tissue Gauges (microtissues) allowed for relatively high-throughput assessment of the rapid dynamics and force generation during cell contraction ^34^. In their model, cells are cultured within a matrix composed of collagen and form around pairs of flexible vertical cantilevers into an array of dense, organized structures comparable to *ex vivo* tissue. High-throughput tensile force measurements can then be calculated from the visible deflection of the cantilevers. More recently, investigators fixed a magnetic microsphere to one of the cantilevers in each microtissue well, and with magnetic tweezers stretched one microtissue at a time for quasi-static stiffness measurements ^35,36^. The limitations in experimental throughput and actuation range of magnetically driven devices were addressed by our recently published Microtissue Vacuum-Actuated Stretcher (MVAS) ^37^. In that publication, the MVAS allowed for high-throughput visualization of cellular remodeling during stretching due to a mostly planar deformation and following chronic (several days) conditioning.

We now present a new microtissue stretcher, the MVAS-force, which enables measurements of tensile force and dynamic mechanical analysis. In contrast to our previous design, only one of the cantilevers in the MVAS-force is actuated through a regulated vacuum pressure while forces are measured simultaneously from the passive bending in the other cantilever. In this article, this new approach allowed us to assess the mechanical properties of microtissues during dynamic loading and upon loading cessation, and to link the changes in mechanics to sub-cellular remodeling using responses to pharmacological treatments and by directly imaging the cytoskeleton. The findings that can be gained from our approach on a cell’s ability, or impaired ability to sense mechanical forces are critical to understand pathways of development, normal function and disease progression in the body.

## Results

### Microtissue Morphology

Within the MVAS-Force, 3T3 fibroblast cells self-assembled around the cantilevers into dense, highly organized, three-dimensional constructs that morphologically resemble tissue. Top-down and cross sectional views of a fully compacted, representative microtissue are shown in fig. 1c. As been shown previously ^34,37,38^, the cells compacted the collagen matrix away from the bottom and sides of the well into a tissue freely suspended around the tops of the cantilevers. The average microtissue thickness measured at its center after four days was 97 ± 2 μm (n=5) and was qualitatively uniform along the longitudinal axis.

**Fig. 1:**
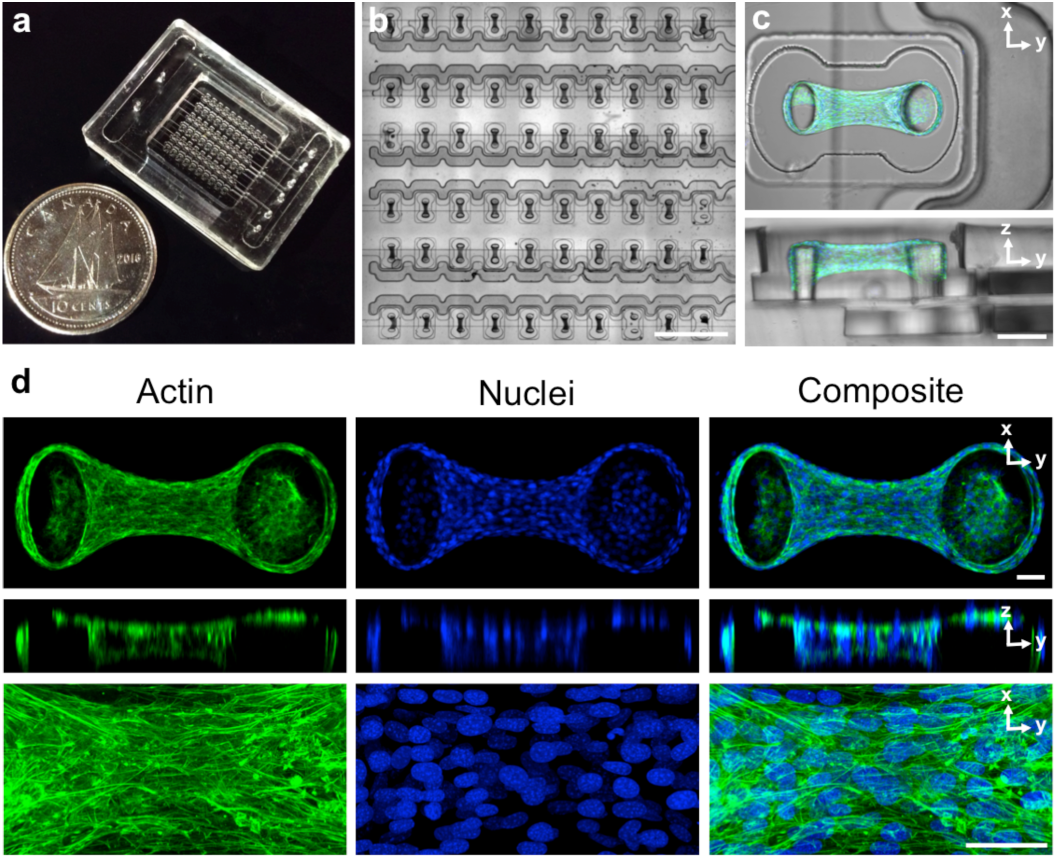
The MVAS-Force allows high throughput dynamic mechanical measurements of 3D cell cultures. The MVAS-force was microfabricated from three photolithographic masters (a). It comprises of an array of microtissue wells each bordered by a controllable vacuum chamber (b). A top-down and cross-section view of a microtissue is shown in (c). Microtissues are dense, organized, three-dimensional cell cultures that are freely suspended around the cantilevers. Max projections of confocal stacks, orthogonal views and high magnification images are shown in (d). The actin cytoskeleton is in green and the nuclei are in blue. Both the cytoskeleton and nuclei show a high degree of organization, aligning between the cantilevers. Scale bars in b,c and d, represent 1mm, 100μm and 50μm, respectively.

Maximum intensity projections with orthogonal slices and centrally located magnified views of F-actin and cell nuclei within a representative microtissue at four days are shown in fig. 1d. Actin was highly polymerized into dense stress fibers that were oriented with the longitudinal axis of the microtissue. The cell nuclei were also mostly aligned with the microtissue and evenly distributed in three dimensions.

### Microtissues strain soften to maintain their mean tension

It has been widely reported that acute dynamic stretching changes the mechanical properties of cells grown on 2D surfaces; they become softer (decreased elasticity) and more fluid-like (increased phase lag) ^7,11,39^. In that regard, we started by investigating whether or not 3D microtissues composed of 3T3 fibroblasts share these behaviors by assessing their dynamic mechanical properties under progressively larger strains at 0.25Hz.

As expected, microtissue storage stiffness (k’) decreased in a strain-dependent manner (N=22, linear regression: R^2^= 0.97, p<0.001), (fig. 2a). Comparing 9% to 1% strain, the average storage stiffness decreased by 26 ± 2% (repeated measures t-test: p<0.001). However, unlike previously published findings on cells in 2D culture, where softening is accompanied with a more fluid-like behavior (or fluidization) ^7^, the phase lag (δ) of microtissues decreased with the strain amplitude (linear regression: R^2^= 0.95, p<0.01) (fig. 2b), indicating a greater amount of energy stored for a given amount dissipated at higher levels of strains. Therefore in contrast to cells in 2D, fibroblast microtissues become more elastic-like as they soften.

**Fig. 2:**
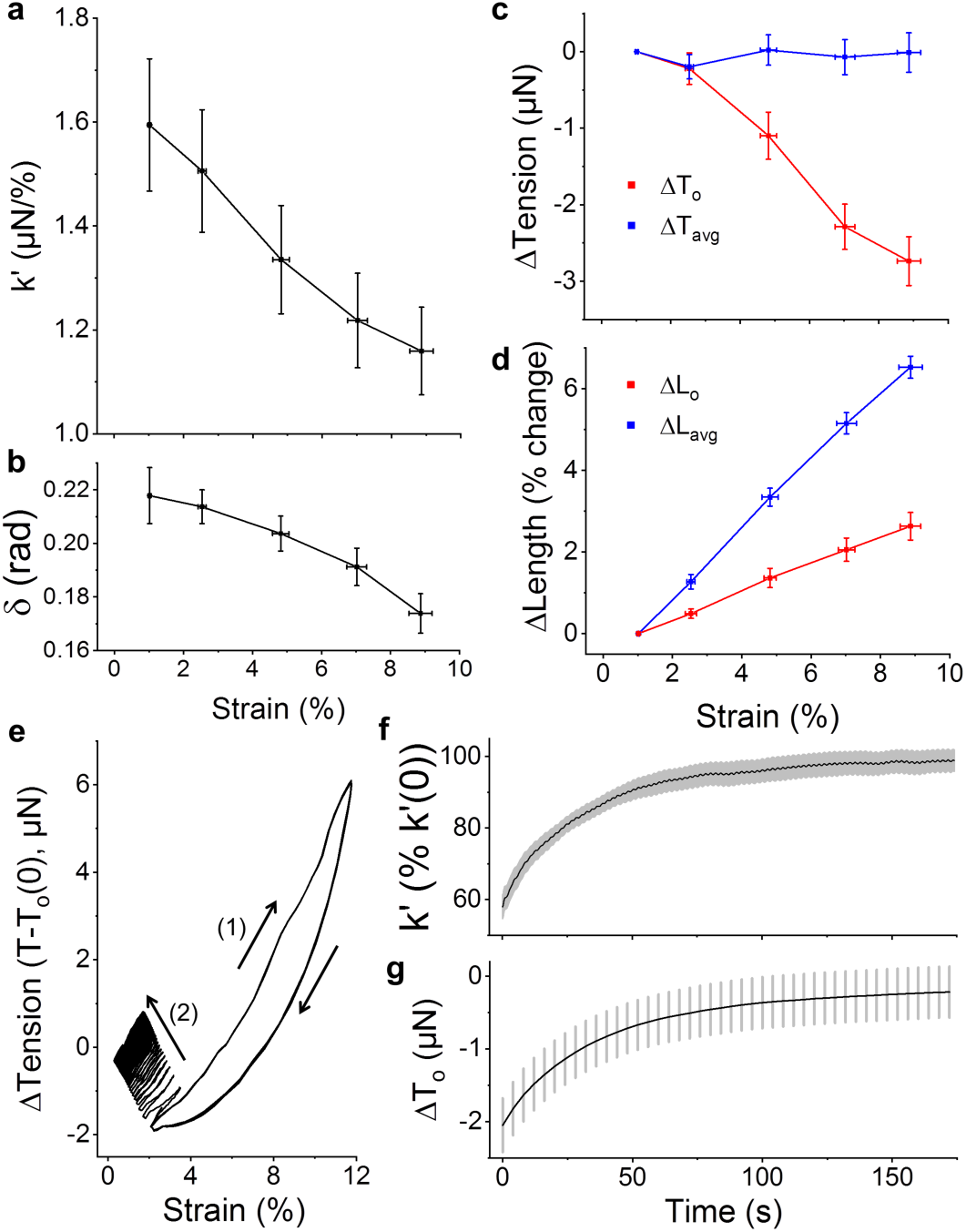
3T3 fibroblast microtissues strain soften to maintain a constant mean stress. The storage stiffness, k’, (a), phase lag between stress and strain, δ, (b) and prestress, T_0_, (c) all decreased with increasing the oscillatory strain amplitude (N=22). Importantly, the decrease to the stiffness and prestress led to a constant mean stress, T_avg_, (c) despite a linear increase to mean microtissue length, L_avg_ (d). Under an auxotonic condition, these behaviors increased the offset microtissue length, L_o_, beyond its initial length. To see whether these responses were reversible, microtissues were oscillated under a large amplitude strain until subsequent loading loops overlapped (1) and then suddenly switched to a small amplitude strain (2) (e). Both the storage stiffness (f) and prestress (g) fully recovered to their initial values over 160 seconds with similar rates (N=8). In this figure, the prestress and mean stress are expressed as the difference from the smallest strain amplitude.

The softening response in microtissues was also accompanied with a decrease to their prestress (T_o_) (linear regression: R^2^=0.97, p<0.001) (fig. 2c). Comparing 9% strain to 1% strain, the average prestress decreased by 2.7 ± 0.3μN (N=22, repeated measures t-test: p<0.001).

Together the softening response and the decrease to the prestress generated a mean tension that was invariant with respect to the strain amplitude (linear regression: p>0.5) (fig. 2c). This behavior occurred despite a linear increase to the mean microtissue length (L_avg_) (linear regression: R^2^=0.99, p<0.001) (fig. 2d) and is indicative of an intrinsic tensional homeostatic response that cells and tissues possess ^29,30^. Moreover, since we considered an auxotonic condition, the minimum microtissue length during a loading cycle (L_o_, ie. the offset length) increased with the stretch amplitude (linear regression: R^2^=0.99, p<0.001) (fig. 2d). Thus, the tension regulating ability of microtissues under dynamic stretching resulted in their lengthening.

The strain softening response was reversible upon returning to small amplitude oscillations (fig. 2e). The stiffness and prestress recoveries are shown in fig. 2f,g, respectively (N=8). After 160 seconds, microtissue stiffness recovered to 99± 3% of its initial value and the prestress agreed well with initial measurements. The recovery curves followed stretched exponential functions (equation 1) with agreeing time (τ) (38 ± 1 vs. 35 ± 6 seconds) and power law constants (β) (0.87 ± 0.01 vs. 0.92 ± 0.08) (SI 1). The offset microtissue length recovered as well with the same dynamics (data not shown).

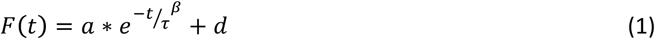

To determine how the softening behavior developed within microtissues during cyclic loading, the first couple of conditioning cycles from a static state were examined in figure 3. At a loading frequency of 0.25 Hz, the mechanical behavior of microtissues adapted over these initial loading cycles (fig. 3a). In that regard, the prestress decreased towards a new set value (fig. 3c), while the mean tension promptly increased and then stabilized back to the resting tension (fig. 3d). As we considered an auxotonic load, these changes resulted in progressive tissue lengthening (fig. 3e). In contrast, when the loading period (400 seconds) was much slower than the recovery time constant (1/f>>τ), the degree of the mechanical adaptation was reduced and predominately occurred over the first loading cycle (fig. 3b-e). Furthermore, when oscillated around an offset strain at 0.0025 Hz, the mean tension did not stabilize to its resting tension (fig. 3d). Therefore tensional homeostasis does not occur when the dynamic loading rate is slower than the rate of tension recovery.

**Fig. 3:**
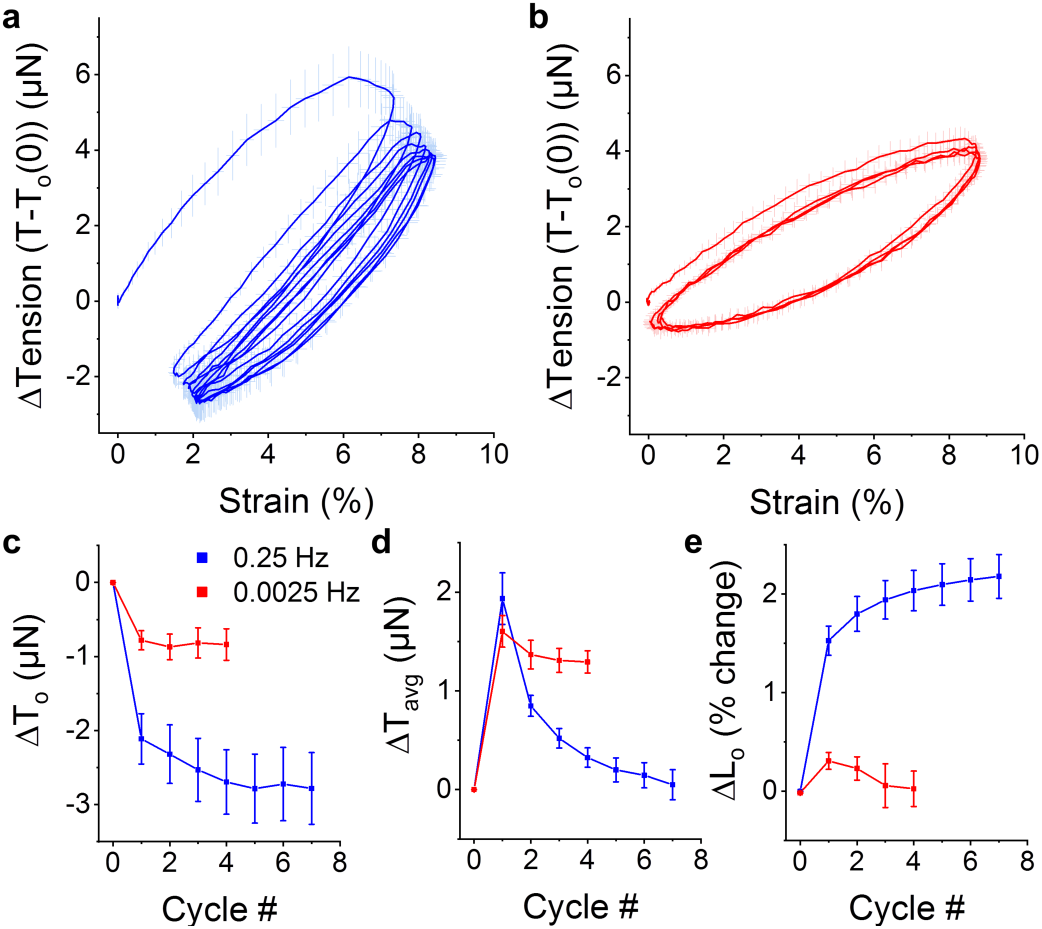
Adaptation to oscillatory loading depends upon the loading frequency. Average (N=6) conditioning cycles from rest at 0.25Hz and 0.0025Hz are shown in (a) and (b), respectively. In both cases, the prestress decreased with subsequent loading cycles towards a new set point but to a greater extent with faster loading frequencies (c). While the mean tension remained elevated when oscillations were slowly applied, it decreased with additional cycles towards the resting tension at 0.25Hz (d). Under an auxotonic condition and when the oscillation frequency was faster than the recovery time constant (1/f≪τ), these behaviors resulted in a progressive lengthening response (e). However, this lengthening response was absent when the oscillation frequency was slower than the recovery time constant (1/f≫τ).

In this section, we have shown that, as with cells in 2D culture, 3D microtissue cultures strain soften to homeostatically maintain their mean tension. Depolymerization of actin filaments ^7,9–11^ and perturbing of myosin motor binding ^12,13,40–42^ are previously hypothesized mechanisms of this response in cells in 2D culture. On the other hand, the involvement of microtubules has been largely overlooked despite their contributions to overall cell mechanics ^43–46^ and their dynamic instability ^47^. Accordingly, we investigated the roles of these three cytoskeletal proteins to microtissue strain softening. We begin with examining the involvement of actin microfilaments.

### Dynamic stretch remodels and depolymerizes actin

To assess the role of the actin cytoskeleton in strain softening, f-actin was depolymerized with Cytochalasin D (CytoD). As expected, CytoD treatment reduced the resting tension, stiffness and phase lag (fig. 4a and SI 2). Importantly, CytoD treatment also muted the softening response (N=16 repeated measures t-test, p<0.001) (fig. 4b). In fact the stiffness of CytoD treated microtissues was independent from the strain amplitude (repeated measures t-test, P>0.05). Upon stretch cessation, CytoD treatment also prevented tension recovery; further demonstrating that CytoD treated microtissues do not soften (fig. 4c). These results indicate that strain softening is dependent upon changes to a densely polymerized actin cytoskeleton.

**Fig. 4:**
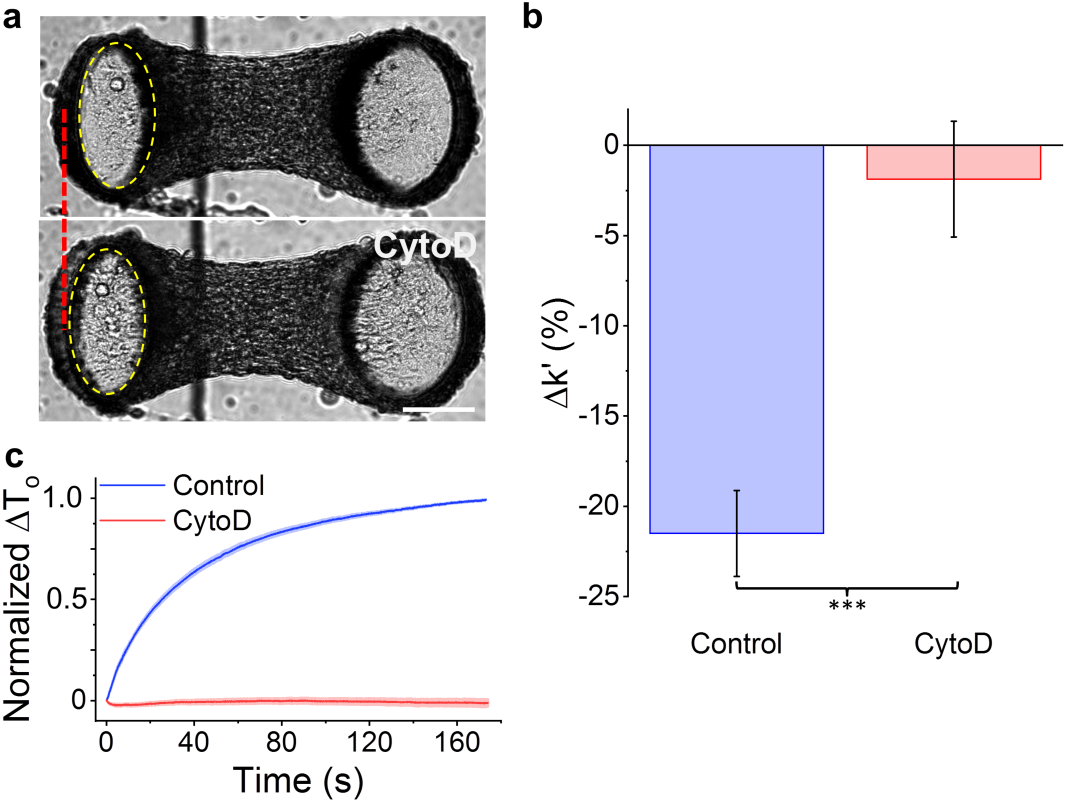
Softening requires an intact actin cytoskeleton. Images of a microtissue prior to and following CytoD treatment are in panel (a). As shown by the dotted red line and the dotted yellow ellipse that outlines the top of the force-sensing cantilever (left), depolymerization of F-actin with Cytochalasin D (CytoD) visibly moved the cantilever outward, indicating a lower resting tension. Importantly, CytoD treatment reduced the stiffness change under large vs. small amplitude stretching (ie. the amount of strain softening) (b). There was also no tension recovery following stretch cessation (c). The scale bar in (a) represents 100μm.

We have shown that f-actin is required for strain softening in microtissues. To further link the softening response to changes in the cytoskeleton, we labeled actin filaments with a live-cell stain. Then by comparing images taken before and following various durations of static resting or oscillatory stretching, we assessed whether stretching increases the rate of actin remodeling. Heat maps of correlation coefficients showing actin remodeling (areas with low correlation coefficients indicate high remodeling) within a centrally located region of the microtissue are in fig. 5a. Actin remodeling was spatially heterogeneous throughout the microtissue and increased with time under both static and loading conditions. Importantly, compared to the static condition, oscillatory loading significantly increased remodeling of actin filaments (decreased the average correlation coefficient) after one and five minutes (N=6, repeated measures t-test, P<0.05) (fig. 5b).

**Fig. 5:**
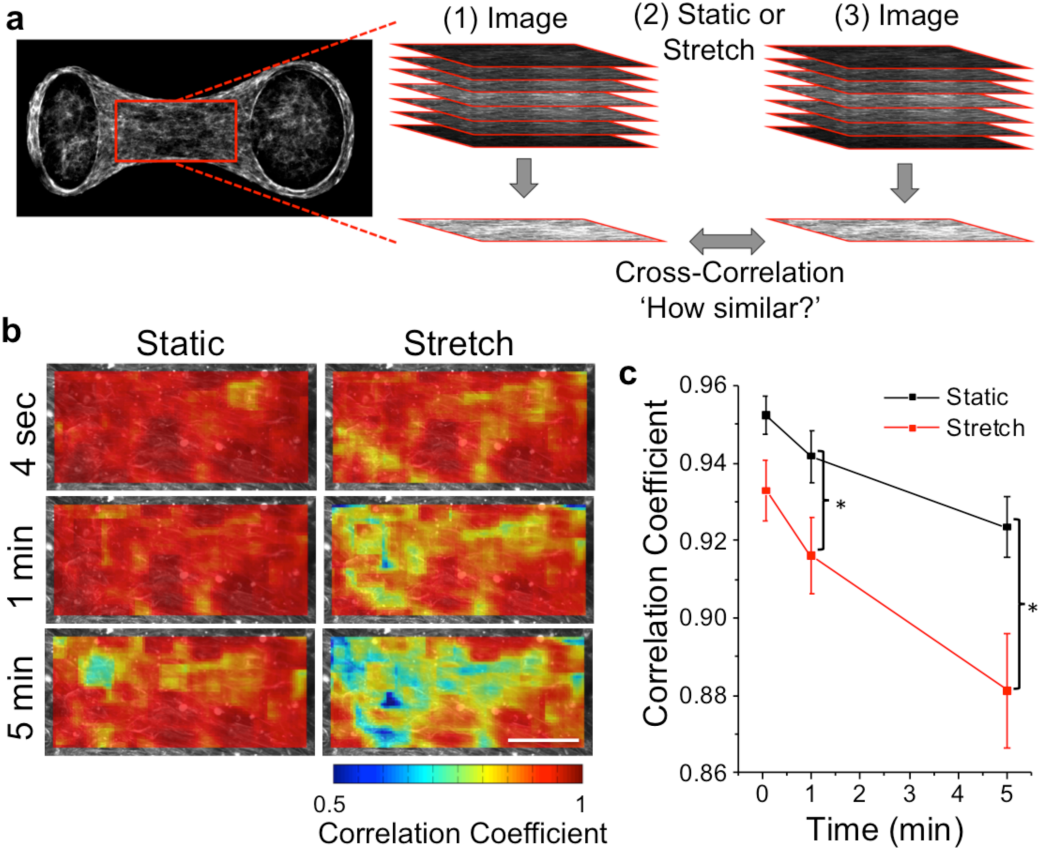
Oscillatory stretch increases remodeling of actin filaments in living cells in 3D culture. The effect of oscillatory stretch on the actin remodeling rate was measured across centrally located regions (212×106µm; red rectangle) in living microtissues using live-cell staining and comparing confocal stacks taken immediately before and after various durations of stretching or static culture (a). Representative heat maps of cross-correlation coefficients show that actin remodeling was spatially heterogeneous and increased with large amplitude stretching vs. static conditions (b). The average correlation coefficient was significantly reduced (ie. a greater amount of remodeling had occurred) when stretching vs. static after 1min and 5min (b). The scale bar in (b) represents 50μm. (*P<0.05; N=6 repeated measures t-test)

To investigate whether remodeling arose simply from organizational changes or depolymerization/repolymerization of filaments, microtissues were fixed and stained immediately following various durations of stretching at 9% strain. Representative images, average heat maps and average f-actin expression per cell (fig. 6a-c, respectively) all indicated that f-actin rapidly depolymerized with oscillatory stretching (N>14). Then to show that f-actin also repolymerizes following stretch cessation, microtissues were fixed and stained after various durations of recovery following five minutes of stretching. Average heat maps and f-actin expression per cell (fig. 6b,d) show complete recovery to initial expression values (t-test P>0.05) (N>11). Although our time resolution of f-actin expression was poor and the uncertainties are large, the rate of f-actin recovery appeared to be within the same order of magnitude (tens of seconds) as the rate of tension and stiffness recovery, suggesting that the mechanical measurements reflect actin repolymerization.

**Fig. 6:**
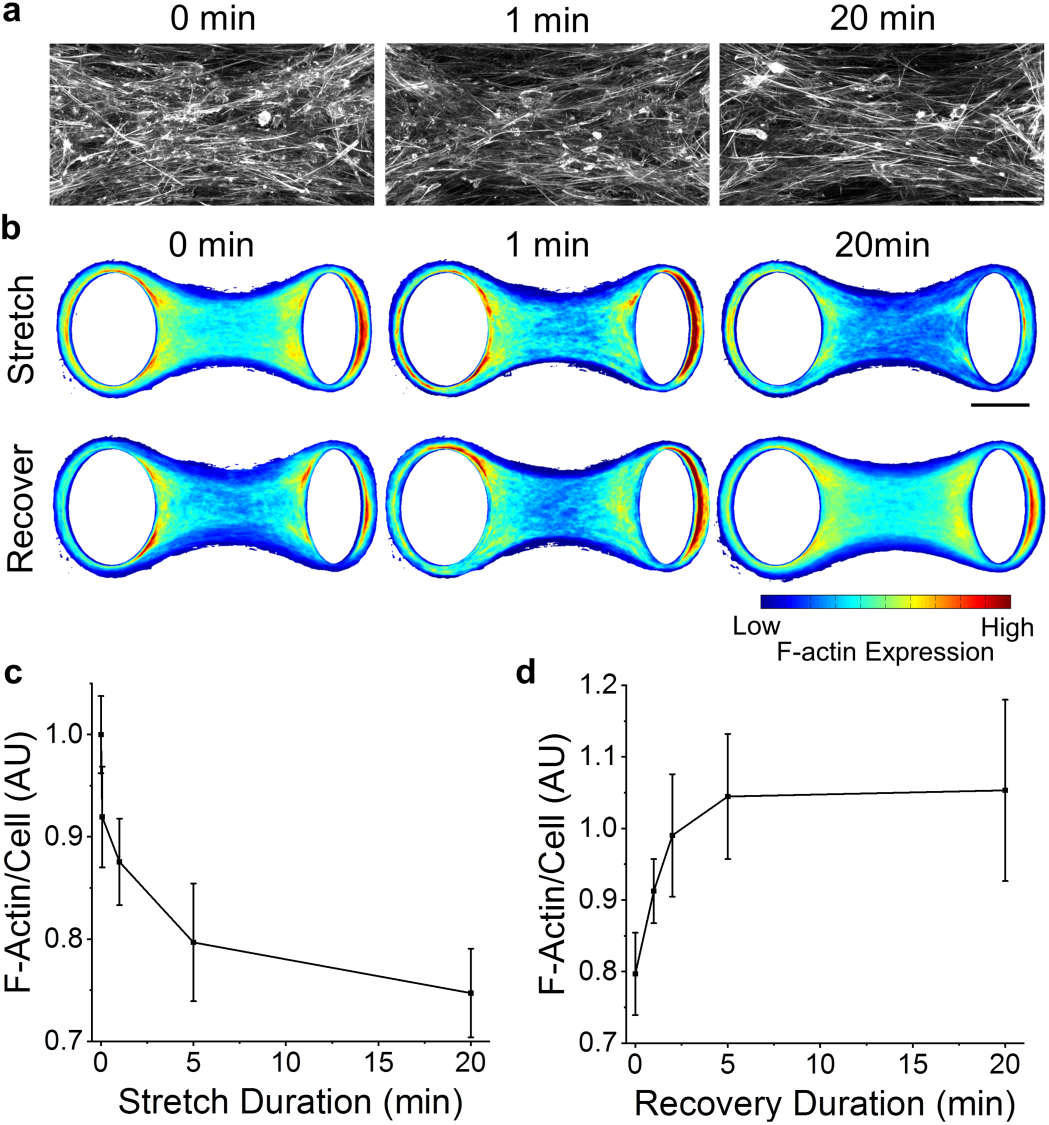
F-actin depolymerizes with stretching and repolymerizes upon stretch cessation. Representative images after different durations of stretching show that there were fewer actin filaments with longer stretch durations (a). F-actin expression in average heat maps was similarly reduced with stretch duration (b) (N>14). Moreover, f-actin expression recovered to initial values upon stretch cessation (N>11). The average actin expression normalized to the number of cells under various durations of stretching and recovery are shown in (c) and (d), respectively. The scale bars in (a) and (b) represent 50 and 100μm, respectively.

### Myosin and microtubules do not contribute to strain softening

We have identified that actin filaments play a major role in the strain softening response of 3D microtissues, however, the mechanical behavior of cells ^43–46,48^ and microtissues is also highly dependent upon myosin activity and microtubules (fig. 7a and SI 2). Accordingly, to assess the contribution of myosin and microtubules to strain softening, we examined the response following myosin inhibition with blebbistatin (Bleb) and microtubule depolymerization with nocodazole (Noco).

**Fig. 7:**
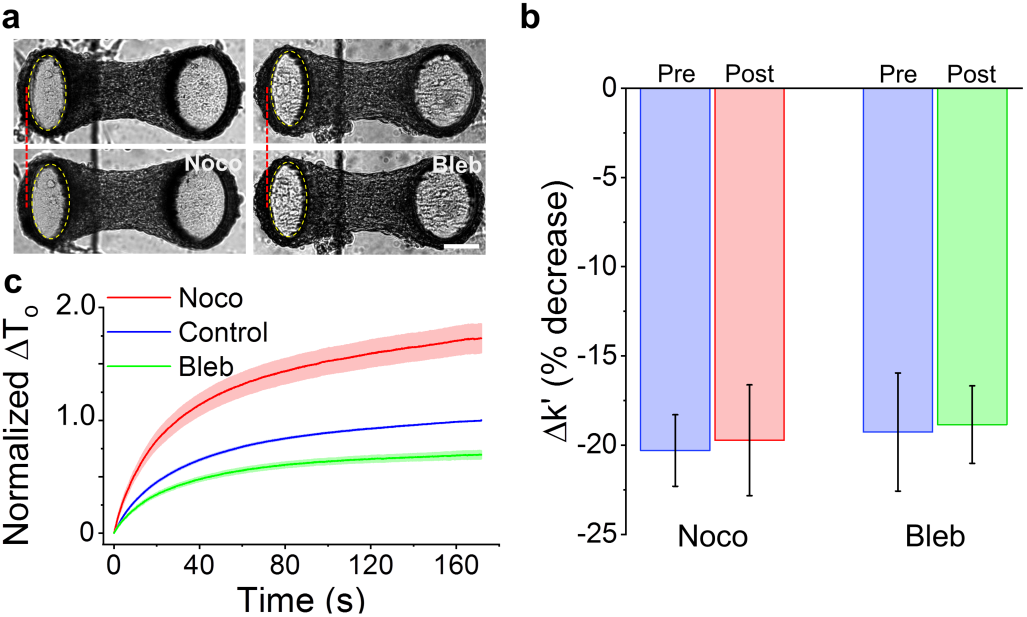
Microtubules and myosin do not contribute to softening. Microtissues prior to and following nocodazole and blebbistatin treatments are shown in (a). Microtubule depolymerization with nocodazole moved the force-sensing cantilever inward, indicating increased prestress. In contrast, myosin-II inhibition with blebbistatin moved the cantilever outward, indicating decreased prestress. Neither treatment changed the amount of strain softening in terms of percent change (b). Furthermore, microtubule depolymerization increased the tension recovery while myosin inhibition decreased recovery (c) but neither treatment changed the time constant of the recovery response. The scale bar in (a) represents 100μm.

Myosin inhibition with blebbistatin decreased microtissue stiffness (N=10, repeated measures t-tests, P<0.05) and prestress (p<0.01) (SI 2). Myosin inhibition, however, did not affect strain softening (N=10, repeated measures t-test, P>0.05). As expected, it did reduce the tension recovery (fig. 7c) because of the decrease in prestress that unsurprisingly accompanied myosin inhibition. However, as there was no change in the rate of recovery, myosin cycling was not likely responsible for the recovery following softening (SI 1). Although it is possible that there was incomplete inhibition of myosin motors, it is unlikely according to blebbistatin’s measured dose-response curve (SI 3). Moreover, even with incomplete inhibition, one would still expect a decrease in the softening response if perturbing of myosin binding were responsible for the strain softening response of microtissues.

In keeping with the hypothesis that microtubules are mainly compressive elements that oppose acto-myosin activity ^43–46^, microtubule depolymerization increased microtissue stiffness (N=15, repeated measures t-tests, P<0.001), and prestress (P<0.001) (SI 2). Microtubule depolymerization, however, had no effect on strain softening in terms of the percent change to the storage stiffness (P>0.05) (fig. 7b). It did increase the absolute tension recovery (fig. 7c) as expected, because of the increase in prestress that accompanied microtubule depolymerization. However, again the molecular mechanism for softening was not likely affected by microtubule depolymerization, as there was no change to the rate of recovery (SI 1). Interestingly, oscillatory stretching did increase remodeling in microtubules with significant differences from static conditions after one and five minutes of stretching (SI 4) (N=7, repeated measures t-tests, p<0.01 and p<0.05, respectively). However, stretching did not change the degree of microtubule polymerization per cell (SI 4) (N>14, 1-way ANOVA).

### Strain softening is a conserved response for microtissues

We have shown that microtissues composed of 3T3 fibroblasts, mimicking connective tissue, strain soften through actin depolymerization. To assess whether this behavior is unique to fibroblasts or is shared among other cell types, we assessed the strain-softening responses in microtissues composed of human airway smooth muscle cells (HASM) or skeletal muscle cells (C2C12). We found that both smooth and skeletal muscle microtissues strain softened (linear regression: R^2^= 0.99, 0.98 P<0.001, 0.001, respectively) (fig. 8a). The phase lag of skeletal muscle microtissues decreased with strain (linear regression: R^2^=0.99, P< 0.001), whereas in smooth muscle microtissues it did not (R^2^=0.43, P=0.14) (fig. 8b).

**Fig. 8:**
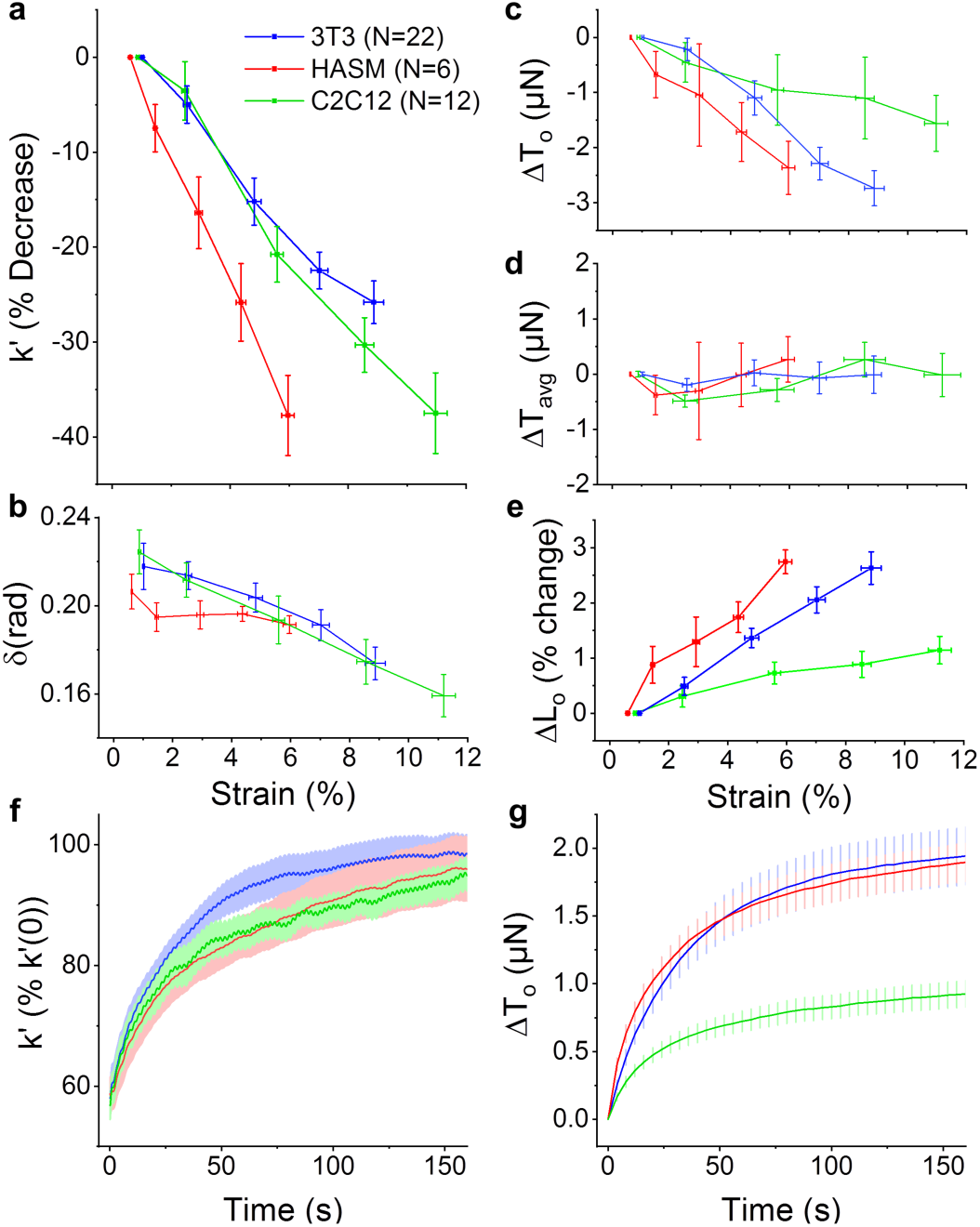
Strain softening is a conserved response in microtissue cultures. Microtissues composed of fibroblast (3T3), human airway smooth muscle (HASM), or skeletal muscle (C2C12) cells all strain softened with similar changes to their stiffness (a), phase lag (b), and prestress (c). For all cell types, the mean tension was invariant with the stretch amplitude (d) despite a linear increase to their mean lengths (data not shown). Together these behaviors led to an increased offset microtissue length (e). 3T3, HASM and C2C12 microtissues also shared similar recovery dynamics upon stretch cessation in terms of the rates of their storage stiffness (f) and prestress (g) recovery.

In addition, as with fibroblast microtissues, HASM and C2C12 microtissues shared strain-dependent decreases to their prestress (linear regression: R^2^= 0.98, 0.94 P<0.001, 0.01, respectively) (fig. 8c). Furthermore, the decrease in their stiffness and prestress likewise led to invariant mean stresses (linear regression: P>0.05) (fig. 8d) despite large changes to their mean lengths (linear regression: both R^2^=0.99, P<0.001, data not shown). Again, these adaptations resulted in microtissue lengthening (ie. L_o_ increased with the strain amplitude) (linear regression: linear regression: both R^2^=0.95, P<0.01) (fig. 8e).

Upon decreasing the strain amplitude, the stiffness and tension of HASM and C2C12 microtissues recovered as previously observed with the fibroblast cultures (fig. 8f,g). The offset microtissue length recovered as well (data not shown). The time constant for stiffness recovery in HASM cells was statistically greater than either 3T3 or C2C12 microtissues (1-way ANOVA, P<0.05) (SI 1). On the other hand, the time constants for tension recovery agreed well with each.

## Discussion

We aimed to assess the response to acute oscillatory stretch in cells grown in conditions that mechanically and biologically recapitulate the 3D environment that a cell would experience were it in the body. In order to fulfill this goal, we developed a novel approach to allow high throughput measurements of dynamic mechanical properties and direct visualization of the cytoskeleton in 3D microtissue cell cultures. Our approach consists of an array of vacuum-driven actuators to stretch microtissues and optically tracked force-sensors to measure their mechanical behavior. Advantages and limitations of our approach are summarized in SI 5. In using our approach, we showed that microtissues soften under dynamic stretching through actin depolymerization. Furthermore, through this softening behavior, tissues homeostatically maintain their mean tension, which results in stretch-induced tissue lengthening. These results are further discussed the sections that follow.

### Microtissues soften under dynamic loading

We showed that the prestress and storage modulus of living 3D microtissue cultures composed of three different cell types decreased with the dynamic strain amplitude. This finding agrees well with previous reports in cells in 2D culture ^7,39,49^, and ex vivo tissue strips ^12,13^. Furthermore, it may reflect an important mechanism through which tissues maintain their lengths, and thereby, homeostasis throughout the body. For example, strain softening may explain how a large tidal stretch from a deep inspiration can open contracted airways in healthy lungs ^12,13^ and could contribute to the regulation of blood pressure in arteries ^15^. Accordingly, under auxotonic conditions, the reported softening behavior of microtissues also led to lengthening. For our approach, the auxotonic resistance is set by the bending stiffness of the force-sensing cantilever. In the body, this resistive force is analogous for example to the parenchymal tethering stiffness in the lung. A loss of these tethering force ^50^ may explain why the normal dilating response from a deep inspiration does not occur in asthmatic airways ^16^.

In addition to softening, ex vivo tissue and cells in 2D have long been reported to exhibit a more fluid-like behavior when stretched ^12,13,51–53^. More recently this fluidization response of cells has been associated with that of a class of inert materials called soft glasses (eg. foams, dense emulsions, pastes and slurries) ^7^. In contrast to these reports and the soft glassy rheology hypothesis, skeletal muscle and fibroblasts microtissues actually became more elastic-like with greater strain amplitudes, and smooth muscle microtissues did not phase transition. This apparent contradiction could arise from environment differences between growing cells in our 3D microtissues and past reports done in 2D culture or differences in the loading protocol. It is also a possibility that the fluidization response was hidden by the elasticity of the ECM. After removing most of the cellular component to the mechanical behavior of the microtissue with CytoD treatment, the remaining behavior, primarily describing the contribution of the matrix, is elastic and contributes little to energy dissipation (SI 2). In turn, a more elastic-like behavior could be perceived as the cells soften (and perhaps fluidize) because the cellular contribution to the overall mechanical behavior becomes less significant compared to the elasticity of the matrix. It would, however, still be of interest in future work to assess whether or not microtissues follow the same time-scale invariance ^54^ that has given traction to the hypothesis that the mechanical behavior of cells follow Sollich’s (1997) theory on soft glassy rheology ^55^.

Upon strain cessation, we showed that microtissue stiffness returned to pre-intervention values along well-conserved trajectories that could be modeled over three magnitudes of time with a stretched exponential. Stretched exponentials have previously been used to describe relaxation processes in disordered systems ^56,57^ and can appear from a linear superposition of simple exponential functions with a nontrivial distribution of relaxation times ^58^. It, therefore, should not be overly surprising that such a function can appear from the complex nature of the cytoskeleton and the added intricacies that arise when considering an aggregate of cells. In spite of this, we found that the recovery time constants for prestress and stiffness were between 35 and 43 seconds for all tested treatments that showed a recovery response and cell types (except for stiffness recovery in HASM microtissues). This time constant appears to be within the same order of magnitude as previously reported recoveries in cortical actin stiffness following transient stretching cells in 2D culture ^7^. Unfortunately the authors of that work did not fit their curves to stretched exponentials. They did, however, concede that recovery occurs with timescales that grow with the elapsed time since stretch cessation and is slower than an exponential process, which are characteristic features of stretched exponentials.

We believe that these abovementioned responses are related to the tensional homeostasis behavior that has been previously reported in 3D cell cultures in response to quasi-static loading ^29,30^. In that regard, it is known that cells will change their internal contractility to oppose external mechanical loads to maintain a set point tension. In keeping with this concept, we found that the decrease in the prestress and stiffness of well-adapted microtissues was sufficient to maintain a constant mean tension across increasingly larger strain amplitudes with increasingly greater mean lengths. Thereby our work extends tensional homeostasis to conditions of dynamically applied loading. In addition we found that the adaptation response occurred at a physiological loading rates, but as one may expected from living matter capable of repair, it was absent when loading was applied slower than the tension and stiffness recovery time constants.

In the literature strain softening exists in a paradox with a large number of studies reporting strain stiffening and actin reinforcement in response to stretch ^59^. Under a sustained stretch or examining different locations along the stress-strain curve, reconstituted crosslinked actin gels ^60,61^, cells ^62^, and 3D cell cultures ^26,63^ have all been shown to strain stiffen. This nonlinear effect arises from reorganization of actin filaments ^60,61^ and, for tissues, reorientation of cells and percolation of these local effects across the cell culture ^63^. Although our loading loops were mostly linear, strain stiffening can be observed to a degree at large strain amplitudes (the total harmonic distortion from nonlinearities increased from 0.041 ± 0.002 at 1% strain to 0.152 ± 0.004 at 9% strain (P<0.0001, repeated measures t-test)). However, this effect was still much smaller than the softening behavior produced by dynamic loading. On the other hand, downstream signaling cascades have been reported to cause mechanical stiffening through actin microfilament reinforcement ^49,64–68^ as a sort of negative feedback to maintain localized mechanical stress ^69,70^. In that regard, microtissues have previously been shown to stiffen following 15 minutes of dynamic stretching ^35,71^ and we have also reported actin reinforcement in microtissues under chronic (2 days) conditioning ^37^. Importantly in those investigations quasi-static stiffness measurements and f-actin expression were evaluated following loading and after a period much greater than the time-scale of stiffness recovery and actin repolymerization we report here. That said, over our relatively short experimental window, we did not observe any differences between initial and fully recovered stiffness, prestress, length and f-actin expression measurements.

### Stretch depolymerizes actin in microtissues

Although strain softening of cells and tissues has been widely reported ^7,12,13,39,49^, the molecular mechanism(s) behind this response remains unclear. Here we investigated the contributions of actin microfilaments, myosin II motors and microtubules.

Firstly, in a cell, the actin cytoskeleton is a filamentous network that gives the cell its shape and opposes tensile forces. In 2D culture, stretching of cells has been reported to depolymerize actin filaments^9–11^. Our results in 3D cultured microtissues agree with those observations. In brief, we found that 1) f-actin was necessary for strain softening and the recovery response; 2) actin remodeling in living cells increased with stretch; 3) short-term stretch lowered f-actin expression; and 4) upon stretch cessation, f-actin expression recovered along the same timescale as tension and stiffness recovery. These findings strongly suggest that strain softening, at least in part, arises from actin depolymerization. Our observed timescale of actin repolymerization following cyclic stretch cessation also agreed well with a previous report that measured actin recovery following a step length change in 3D cell cultures ^31^.

Although the regulation of actin involves multiple proteins and is not fully understood, strain induced actin depolymerization has been recently linked to increased cofilin activity. In that regard, knocking down this actin-severing molecule in cells in 2D culture, reduced softening and their actin filaments remained largely intact following a transient stretch ^72^. In addition, myosin Ib has recently been shown to act as a catch-bound actin depolymerase (its affinity for actin, and thus its stress fiber severing capability, strongly increases with the applied load) ^73^, and thus may also promote the strain softening behavior of cells and tissues. On the other hand, the actin recovery response was perhaps mediated in part by zyxin facilitated stress fiber repair. In that regard, it has been previously shown that zyxin localizes to sites of stress fiber fragmentation, and knockdown of zyxin reduces the recovery of contractile force in single cells and leads to more rapid dilation in precision cut lung slices following stretch ^74^. Assessing these proteins in our MVAS-force device may further our understanding of their contributions to the dynamic regulation of the cytoskeleton and the mechanical properties of tissues, and thus, should be a focus of future investigations.

Next, myosin II motors regulate the mechanical behavior of cells by generating tension through crosslinking and actively pulling on actin filaments. Furthermore, strain softening in reconstituted actin-myosin networks has been attributed to the disruption of myosin crosslinks ^40–42^. Perturbing of the binding of myosin has also been implicated in the softening response in airway tissue strips ^12,13^. In contrast, we found that softening in microtissues was invariant on myosin activity and that there was no change to the rate of the recovery response following strain cessation. This strongly suggests that myosin has no role in the softening response of cells in 3D culture.

Lastly, although our understanding of the role of microtubules in cell mechanics is still being refined ^75^, it is thought that they act as compressive struts that oppose actin-myosin contractility, as in tensegrity architecture ^43–46^. Accordingly, and in keeping with several other investigations measuring cell traction forces and stiffness in 2D ^43,76^, we found that depolymerization of microtubules with nocodazole increased microtissue stiffness and prestress (SI 2). Comparably fewer studies have assessed how stretching cells and tissues affects microtubule remodeling and polymerization. That said, microtubules have visually been seen to buckled while cells deform ^45^, and in axons grown in 2D culture, microtubules have been shown to disassemble under large (75%) dynamic loading ^77^. In contrast, we did not observe any changes to microtubule polymerization and nocodazole treatment had no affect on the softening response. Dynamic stretching did, however, increase microtubule remodeling. Whether stretching directly caused disassembly/assembly of microtubules or they simply remodeled in accordance with actin depolymerization, and whether or not the observed microtubule remodeling contributes to changes in the mechanical behavior of cells are interesting questions for future investigations.

## Conclusions

In this article, we presented a new high-throughput approach for assessing both dynamic cell mechanics and for visualization of remodeling at the sub-cellular level in response to stretch within physiologically relevant 3D microtissue cultures. Our approach offers the ability to link behaviors observed in 2D culture to cells within a soft 3D matrix comparable to human tissue, and to connect visual remodeling of the cytoskeleton to changes in mechanical properties. In that regard, we found that fibroblast, smooth muscle, and skeletal microtissue cultures all shared a conserved softening response when dynamically stretched and recovery following stretch cessation. Microtissues responded to maintain their mean stress, and under an auxotonic condition, their response led to lengthening. Furthermore, by directly quantifying cytoskeletal remodeling, these behaviors appeared to arise from rapid actin depolymerization. This suggests that actin microfilaments are sensors of mechanical stretch in cells, and in turn, form a feedback loop to control the mechanical behavior of tissues. The ability of cells to feel and react to mechanical stimuli from their environment is an important mechanism for maintaining homeostasis in the body and is a critical aspect to a full understanding of many pathological disorders.

## Methods

### Device design

Our original MVAS device ^37^ was modified to allow *in situ* measurements of microtissue tension and dynamic stiffness. The MVAS-Force consists of six independently controllable rows of ten microtissue wells (fig. 1). Vacuum chambers border one side of each row of wells. Within each well, there are two cantilevers spaced apart by 500µm. One cantilever is secured on a flexible membrane that deforms to stretch the microtissue when a vacuum is applied through an external electronic regulator (SMC ITV0010) controlled via Labview software (movie 1). The other cantilever acts as a passive force sensor. Its deflection is optically tracked and converted into a force measurement using a spring constant of k_cantilever_=0.834N/m estimated with Euler-Bernoulli beam theory and verified with atomic force microscopy (SI 6).

### Device Fabrication

Devices were fabricated as previously described ^37^ with slight modifications and detailed assembly steps are illustrated in SI 7. Briefly, the MVAS-Force consists of three layers fabricated through mold replication from SU-8 (Microchem) masters made with standard photolithographic methods on polished silicon wafers (Universitywafers.com). All photomasks were ordered from CAD Art Services Inc. The top layer of the MVAS-Force comprises the open-top microtissue wells and enclosed vacuum chambers. The thin middle membrane is fabricated with the cantilevers around which the microtissues compact. Finally the bottom layer contains vacuum chambers that match the top layer, and bottom chambers that equalize the pressure on either side of the membrane to minimize out of plane motion. All three layers were cast in polydimethylsiloxane (PDMS) from the SU-8 negatives with a 10:1 monomer to curing agent ratio and then plasma bonded together. To aid in tracking the bottom of the cantilevers, the middle membrane was fluorescently dyed with Rhodamine B (RhoB).

### Cell culture

NIH3T3 fibroblast (ATCC) and C2C12 skeletal muscle (ATCC) cells were cultured in Dulbecco’s Modified Eagle’s Medium (DMEM) (Hyclone Laboratories Inc.). Human ASM cells (Donor 12) (previously characterized by Gosens *et al*. ^78^) immortalized by stable transfection with human telomerase reverse transcriptase were obtained as a generous gift from Dr. William Gerthoffer (University of South Alabama) and maintained in DMEM/F12 (Invitrogen 11330). All culture media was supplemented with 10% fetal bovine serum (FBS), 100mg/ml streptomycin and 100U/ml penicillin antibiotics (all from Hyclone Laboratories Inc.). Cells were grown at 37°C with 5% CO_2_ on 100mm tissue culture dishes (Fisher) until 80-90% confluent.

### Microtissue fabrication

Microtissue fabrication was performed as described previously ^34,38^, with modifications. Briefly, the device was sterilized with 70% ethanol, and treated with 0.2% Pluronic F-127 (P6866, Invitrogen) for two minutes to reduce cell adhesion. 250,000 cells were resuspended in 1.5mg/ml rat tail collagen type I (354249, Corning) solution containing 1x DMEM (SH30003.02, Hyclone), 44 mM NaHCO_3_, 15 mM d-ribose (R9629, Sigma Aldrich), 1% FBS and 1 M NaOH to achieve a final pH of 7.0-7.4. The cell-collagen solution was pipetted into the MVAS-Force and centrifuged to load ∼650 cells into each well. The excess collagen was removed and the device was transferred into the incubator for 15 minutes to initiate collagen polymerization. An additional 50,000 cells were added and allowed to adhere to the top of the tissues. Excess cells were washed off. Cell culture media was added and changed every 24 hours.

### Force measurements

Microtissue mechanics were deduced from the visible deflection of the force-sensing cantilever while under dynamic loading at 0.25Hz (unless stated elsewise) (movie 1). Prior to measurements, microtissues were preconditioned until subsequent loading loops were superimposable. All measurements were completed at 37°C and 5% CO_2_.

To track cantilever deflection, images of both the tops and bottoms of the cantilevers were captured at 15 frames per second for one minute. The bottom positions were measured by finding their centroids on thresholded fluorescent images of the Rho-B dyed cantilevers. The top of each cantilever was tracked using pattern matching with adaptive template learning in Labview on brightfield images. The deflection of the force sensor was calculated from the difference in the top and bottom positions after accounting for the phase lag caused by the camera delay between the top and bottom images. The deflection was then converted into a force measurement using the cantilever spring constant, k_cantilever_.

Microtissue strain, ε, was defined as the percent change in the length between the innermost edges of the tops of the cantilevers (equation 2):

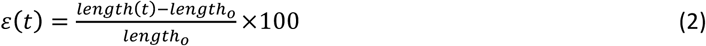

The phase lag, δ, between force and strain was defined as the difference in the phase angles, Φ, at the oscillatory frequency (equation 3):

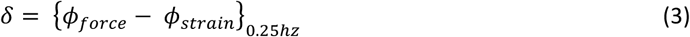

The storage, k’, microtissue stiffness was defined as the ratio of the magnitudes of the Fourier Transforms of force and strain at the oscillatory frequency multiplied by the cosine of the phase lag between force and strain (equations 4):

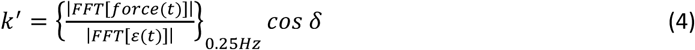

k’ describes the amount of energy that is elastically stored for a given deformation, and δ describes the ratio of energy dissipated to energy stored where in purely elastic samples tan(δ)=0 and in purely viscous samples tan(δ)=inf.

The tension offset or prestress, T_o_, was defined as the magnitude of the Fourier transform of the microtissue force at 0Hz minus the half of the peak-to-peak magnitude of the Fourier transform at 0.25Hz (equation 5).

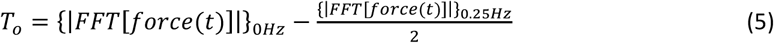

The noise floor for calculating microtissue mechanics and measurements of an elastic standard with our approach are given in SI 6.

To assess the response of microtissue mechanics to stretch, measurements were taken at progressively higher strains. After completing measurements at the largest strain, the recovery response was measured by promptly decreasing the strain amplitude. Stiffness recovery was measured by performing Fourier transforms on intervals spanning three loading cycles.

To assess the role of individual cytoskeletal proteins in contributing to the mechanical properties of microtissues and the strain softening behavior, measurements were taken following 20 minute incubations with either 10µM nocodazole (Noco), a microtubule polymerization inhibitor, 5µM blebbistatin (Bleb), a myosin-II inhibitor, or 10 μM cytochalasin D (CytoD), an actin polymerization inhibitor. As mechanical properties can vary between microtissues, each microtissue was compared to its own pre-treatment value where indicated. To prevent crossover in responses from multiple drugs, only a single treatment was administrated to a microtissue.

### Quantification of cytoskeletal remodeling and polymerization

Images were acquired on a TiE A1-R laser scanning confocal microscope (LSCM) (Nikon) with standard LSCM configurations using appropriate laser lines and filter blocks.

To assess actin and microtubule remodeling in living microtissues in response to stretch, cells were loaded with either 0.1μM SiR-actin or SiR-tubulin with 1μM verapamil 6-12 hours before imaging. Z-stacks were taken before and following 4 seconds (one stretch), one minute and five minutes of static resting and then ∼9% stretching at 0.25hz. Imaging was completed with a 60x 1.2NA water immersion objective to give a centrally located field of view of 212×106µm (1024×512 pixels). Z-stacks were flattened by integrating slices, divided into sub images with a size of 100×100 pixels with 10-pixel spacing, and compared with cross-correlation. The correlation coefficient is a measure of how closely images matched before and after a given condition, and thus is inversely proportional to the amount of remodeling (ie. a low correlation coefficient corresponds to a high degree of remodeling).

To assess f-actin expression and microtubule polymerization, microtissues were fixed *in situ* with 3.5% paraformaldehyde for 15 minutes and permeabilized with 0.5% Triton-X for 5 minutes. Microtissues were left in blocking buffer (5% FBS in PBS) for 40 minutes. Microtubules were labeled with α-tubulin primary antibody produced in mouse (Sigma, T6074) and a rabbit anti-mouse IgG secondary antibody conjugated to Alexa Fluor 488 (Invitrogen, A11059). The actin cytoskeleton was stained with Alexa Fluor 546 Phalloidin (Fisher, A22283), and the nuclei were stained with DAPI (Fisher, D1306). To quantify f-actin expression per cell, Z-stacks were flattened by integration, averaged and normalized to DAPI fluorescence. Microtubule polymerization was quantified with the same method except images were first thresholded to remove any signal from nonpolymerized tubulin.

### Data analysis and statistics

All numerical data are presented as mean ± standard error unless indicated otherwise. Statistical tests, as described in the results, were performed using Originlab 8.5 (Northampton, MA), with p<0.05 considered statistically significant.

## Acknowledgements

M.W. is supported by OGS (Ontario Graduate Scholarship). The authors acknowledge support from individual NSERC Discovery Grants (M.G. and A.E.P.). M.G. acknowledges the Canadian Foundation for Innovation. A.E.P also acknowledges generous support from the Canada Research Chairs program.

## Additional Information

### Author Contributions

M.W. performed data acquisition and analysis and wrote the manuscript. P.R. contributed to data acquisition. All authors contributed to the study design and revised the manuscript.

### Data Availability

The data generated during the current study is available from the corresponding author upon reasonable request.

