## Supplementary Information for "Structural and mechanical remodeling of the cytoskeleton maintains tensional homeostasis in 3D microtissues under acute dynamic stretch"

*Condensed Running Title: The cytoskeleton in microtissues*

Matthew Walker[1], Pauline Rizzuto[2], Michel Godin[3,4,5], Andrew E. Pelling[1,3,6,7]*

[1]Department of Biology, Gendron Hall, 30 Marie Curie, University of Ottawa, Ottawa, ON, K1N5N5 Canada
[2]Université Côte d'Azur, 28 Avenue de Valrose, Nice, 06108 France
[3]Department of Physics, 150 Louis Pasteur pvt., University of Ottawa, Ottawa, ON K1N 6N5 Canada
[4]Department of Mechanical Engineering, Colonel By Hall, 161 Louis Pasteur, University of Ottawa, Ottawa, ON K1N6N5 Canada
[5]Ottawa-Carleton Institute for Biomedical Engineering, Colonel By Hall, 161 Louis Pasteur, University of Ottawa, Ottawa, ON K1N6N5 Canada
[6]Institute for Science Society and Policy, Simard Hall, 60 University, University of Ottawa, Ottawa, ON, K1N5N5 Canada
[7]SymbioticA, School of Anatomy, Physiology and Human Biology, University of Western Australia, Perth, WA, 6009

*Keywords:*
Microtissue, cell mechanics, 3D cell culture, cytoskeleton, actin, microtubules, myosin, microfabrication, lab-on-a-chip, fibroblasts, airway smooth muscle, skeletal muscle

* Author for correspondence
Andrew E. Pelling
598 King Edward
University of Ottawa
Ottawa, ON K1N 6N5
Canada

Web: http://www.pellinglab.net

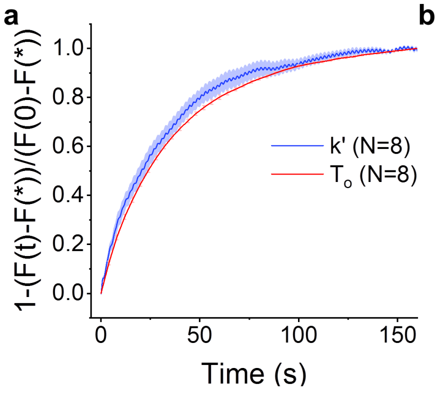


| | a | τ (s) | β | d | r² |
|---|---|---|---|---|---|
| $T_o$ | -1.9 ± 0.2 μN | 38 ± 1 | 0.87 ± 0.01 | 1.9 ± 0.1 μN | 0.9991 ± 0.0003 |
| k' | -0.73 ± 0.08 μN/% | 35 ± 6 | 0.92 ± 0.08 | 1.8 ± 3 μN/% | 0.995 ± 0.002 |

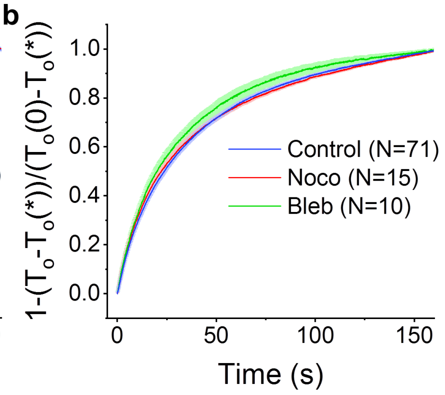


| | a (% Control) | τ (s) | β | d (% Control) | r² |
|---|---|---|---|---|---|
| Control | - | 43 ± 2 | 0.79 ± 0.01 | - | 0.9988 ± 0.0003 |
| Noco | 85 ± 13 *** | 42 ± 3 | 0.71 ± 0.02 ** | 83 ± 13 *** | 0.995 ± 0.002 |
| Bleb | -31 ± 5 *** | 38 ± 7 | 0.79 ± 0.02 | 31 ± 5 *** | 0.9989 ± 0.0003 |

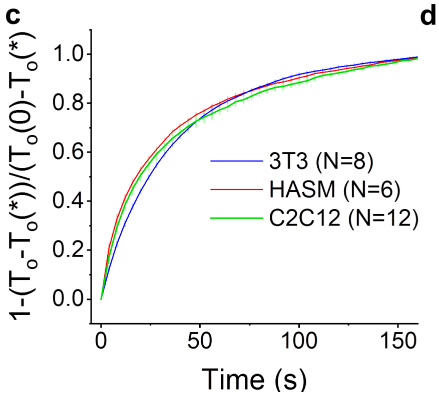


| | a (μN) | τ (s) | β | d (μN) | r² |
|---|---|---|---|---|---|
| 3T3 | -1.9 ± 0.2 | 38 ± 1 | 0.87 ± 0.01 | 1.9 ± 0.1 | 0.9991 ± 0.0003 |
| HASM | -2.0 ± 0.1 | 36 ± 2 | 0.66 ± 0.03 | 2.0 ± 0.1 | 0.9988 ± 0.0004 |
| C2C12 | -1.0 ± 0.1 | 43 ± 6 | 0.67 ± 0.04 | 1.0 ± 0.1 | 0.9971 ± 0.0004 |

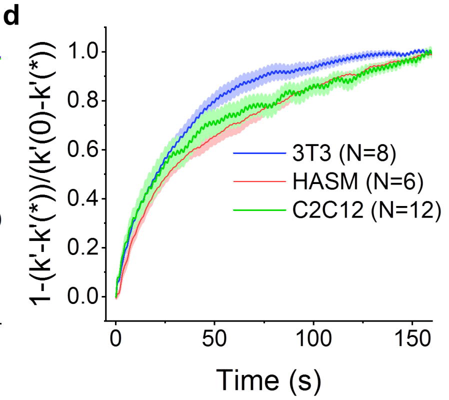


| | a (%) | τ (s) | β | d (%) | r² |
|---|---|---|---|---|---|
| 3T3 | -42 ± 2 | 35 ± 6 | 0.92 ± 0.08 | 100 ± 4 | 0.995 ± 0.002 |
| HASM | -56 ± 12 | 138 ± 48 | 0.7 ± 0.1 | 110 ± 13 | 0.9955 ± 0.0008 |
| C2C12 | -42 ± 3 | 41 ± 7 | 1.0 ± 0.1 | 96 ± 3 | 0.985 ± 0.006 |

**SI 1: Microtissue recovery shares similar dynamics across pharmacological treatments and cell types**. For microtissues composed of 3t3 cells, tension and stiffness recovery followed similar trajectories. They both fit well to stretch exponentials with agreeing time (τ) and power (β) constants (a). Microtubule depolymerization (Noco) or myosin inhibition (Bleb) did not change the time constant and only microtubule depolymerization changed the power law constant (repeated measures t-tests). The tension recoveries for 3T3, HASM, and C2C12 microtissues are shown in (c). The tension recovery in C2C12 microtissues was significantly less than 3T3 ($P<0.01$) and HASM ($P<0.001$) (1-way ANOVA). There was no change in the time constant. The power law constant for 3T3 microtissues was greater ($P<0.001$) than C2C12 or HASM. The stiffness recoveries for 3T3, HASM, and C2C12 microtissues are shown in (d). The time constant for HASM recovery was greater than either 3T3 or C2C12 microtissues ($p<0.05$). There were no other significant differences.

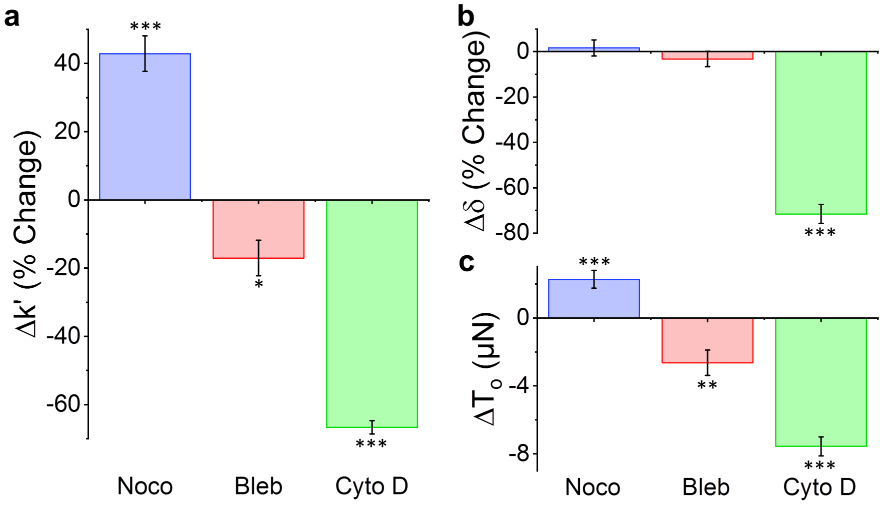


**SI 2: Pharmacological responses.** Microtissue storage stiffness (a), phase lag (b) and prestress (c) change in response to microtubule depolymerization (noco) (N=15), myosin inhibition (bleb) (N=10) and actin depolymeriza5ion (cytoD) (N=16). Microtubule depolymerization significantly increased storage stiffness and prestress. Myosin inhibition or actin depolymerization decreased storage stiffness and prestress. Actin depolymerization also significantly reduced the phase lag. (*P<0.05; **P<0.01; ***P<0.001; repeated measures t-tests).

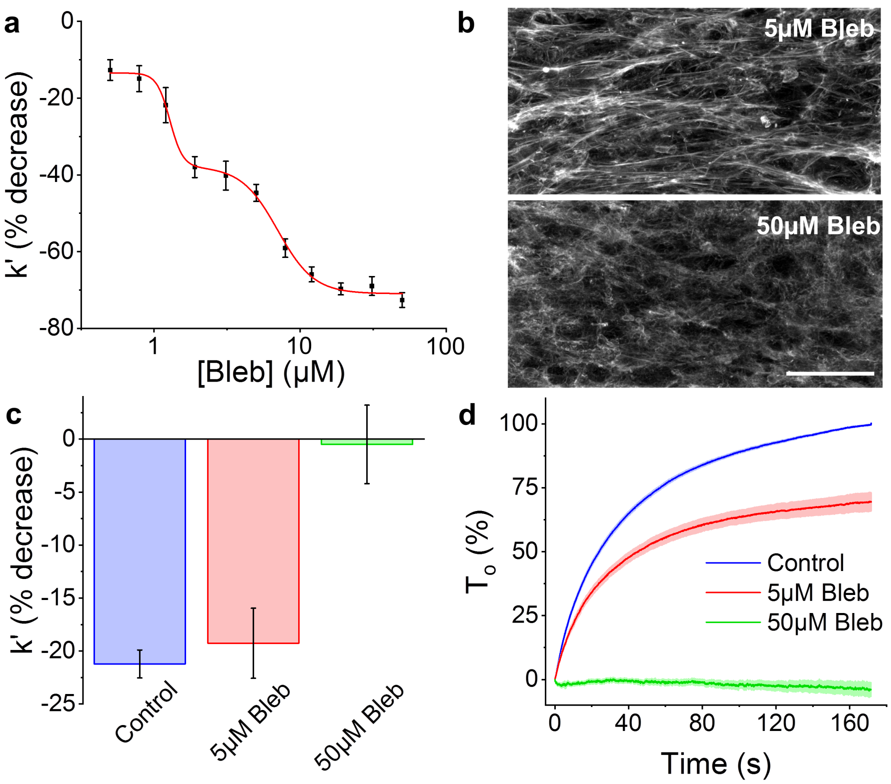


**SI 3. Strain softening does not depend upon myosin activity.** The dose-response curve for blebbistatin is biphasic (a). Biphasic responses result from multiple mechanisms of action. Blebbistatin is a selective myosin-II inhibitor but at higher concentrations other changes to structural proteins may occur. For example following a treatment with 5µM blebbistatin, the f-actin cytoskeleton is largely intact, whereas following a 50µM dose f-actin is more unorganized and not densely polymerized into stress fibers (b). Therefore it is reasonable to suspect that the first plateau of the dose-response curve, which ends roughly at 5µM, represents myosin-II inhibition, while at greater concentrations the response is more inline with actin depolymerization. At 5µM there was no change in softening in terms of percent stiffness change (c) (1-way ANOVA). Only the absolute value of prestress recovery changed (d), which is to be expected because blebbistatin decreases the prestress on its own. On the other hand, at 50µM, and because of its action to depolymerize f-actin, the response reflects changes seen in Fig. 3 with cytochalasin D- no softening and no recovery. The scale bar in (b) represents 50um.

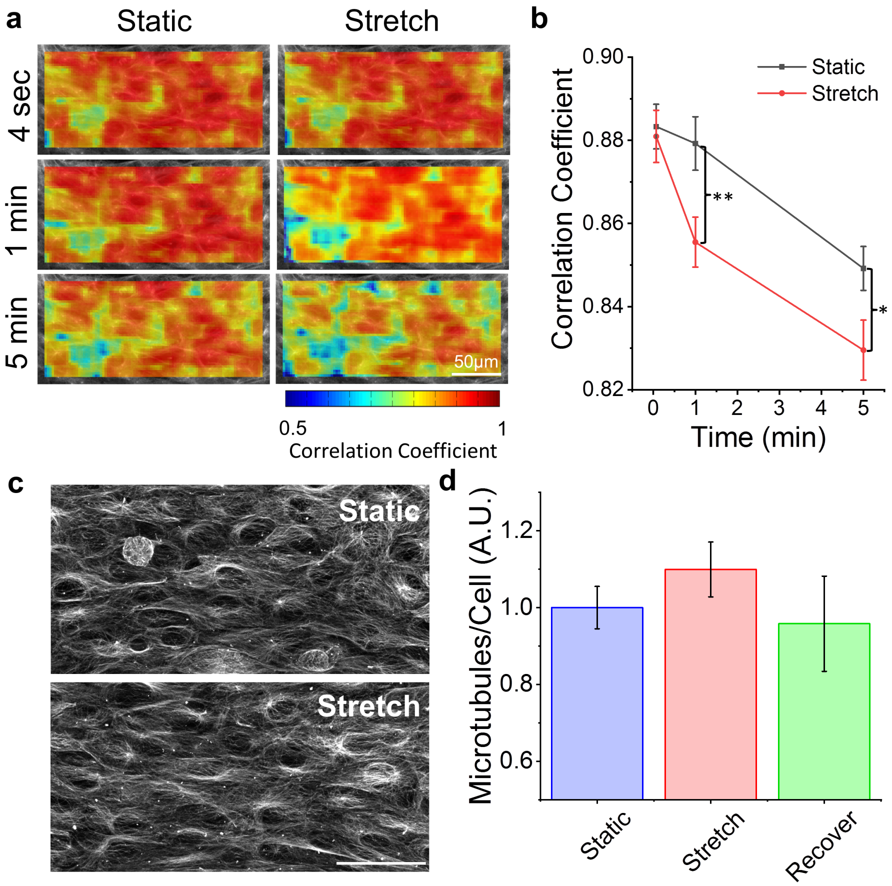


**SI 4: Microtubules remodel under oscillatory stretching but polymerization does not change**. Representative heat maps of microtubule remodeling after various durations under static and stretching conditions are shown in (a). The average correlation coefficients are in (b). At 1min and 5min there was significantly more remodeling in microtubules when under oscillatory loading than in static (* $P<0.05$, ** $P<0.01$; N=7, Paired t-tests). This was not unexpected especially considering we saw similar strain-dependent remodeling in f-actin filaments and microtubule and actin dynamics have been previously linked in the literature.[1] Unlike actin, however, stretching caused no observable change in microtubule distribution/organization (c) and there was no difference in the amount of polymerization per cell after 5min of stretching (N=15) or 5min of recovery (N=14) compared to static (N=16) (d) (1-way ANOVA). The scale bar in (c) represents 50um.

**SI 5: Advantages and limitations of the MVAS-force**

The main advantage of our device over previous methods (ie. atomic force microscopy (AFM), optical magnetic twisting cytometry (OMTC) and optical/magnetic tweezers) used to assess micro-rheology of cells is that in our device cell mechanics are assessed within a 3D physiologically relevant environment, whereas in previous methods, cells are grown on 2D substrates. Although direct mechanical comparisons between cells grown on 2D and in 3D environments are lacking, a third dimension for cell adhesion is known to significantly affect the distribution and structure of the cytoskeleton.[2] In addition to possibly affecting mechanical properties, we are starting to appreciate that the dimensionality of the extracellular environment may influence how cells respond to mechanical stimuli.

Another advantage of our device over previous micro-rheology methods is that the uniaxial tensile deformation used to measure mechanical properties resembles the deformation a cell would experience in airways during inspiration or in blood vessels during systole. In comparison local shear from magnetic beads in OMTC or compressive deformation from an AFM tip have limited relevance. In our device the length-scale of deformation is the size of the cell whereas the deformation during OMTC or with a fine AFM tip is very local and volumetrically relatively small compared to the size of a cell. This difference has important consequences when comparing mechanical properties between microrheology methods. First, the measurement of elasticity can vary over four magnitudes depending upon the length scale of deformation.[3] Secondly, stress dissipation from a volumetrically small deformation may arise solely from frictional stress between cytoskeletal filaments, and thus follows the structural damping law,[4,5] whereas at the length scale of the entire cell, dissipation may arise from both the cytoskeleton and viscous damping through cytosolic fluid movement.[6]

Static measurements of microtissue elasticity have been previously investigated using a magnetic actuation system.[7,8] In this closely related method, a ferrous bead is fixed to one of the cantilevers and moved by magnetic tweezers while the other cantilever is used as a force sensor. Although this method has provided valuable insights into how tissue-level forces are generated from cells, actuation through magnetic tweezers is not suitable for long-term simultaneous conditioning of multiple microtissues and the device fabrication throughput is limited. Our vacuum actuation approach overcomes these disadvantages.

Perhaps one limitation of the MVAS-Force device is that it requires specialized equipment for master fabrication and bonding device layers together. The equipment (plasma machine, mask aligner, and spin coater) is, however, common to microfabrication facilities and presently available on most university campuses.

In this article we reported, for the first time, on cytoskeletal remodeling in response to dynamic stretching in living cells in 3D cell cultures. Although we believe that the ability of our device to allow high resolution imaging of living cells in a physiologically relevant environment marks a significant improvement over previous methods, there is some difficulties that remain. Firstly, despite our best efforts to make a device with minimal out of plane motion, there is a vertical deflection of roughly 0.5μm/% strain. Although this is not a great deal of vertical motion, admittedly it is enough to change focal planes when imaging with high numerical aperture objectives. This was not a problem for the work in this article because all imaging was done at 0% strain. If in future work, images would like to be compared at different strains, this limitation could be overcome by programing a z-stage to work in conjunction with the MVAS-force. Secondly, although microtissues are far easier to image than centimeter scale 3D cultures, imaging does still suffer from fluorescence attenuation and longer acquisition times when capturing z-stacks compared to imaging cells on 2D glass or plastic surfaces. For this reason, imaging cells in 2D culture will likely remain for now the most common method for assessing cytoskeletal remodeling. However, the MVAS-force is a suitable alternative for answering whether or not similar remodeling occurs in a more relevant 3D environment.

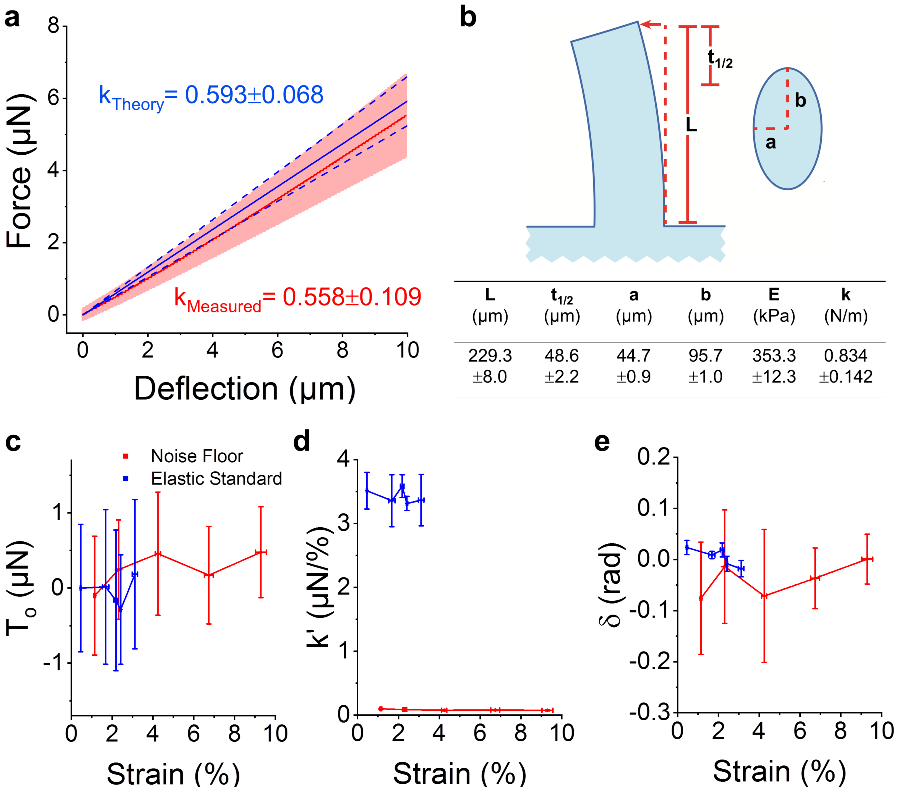


| L (μm) | $t_{1/2}$ (μm) | a (μm) | b (μm) | E (kPa) | k (N/m) |
|---|---|---|---|---|---|
| 229.3 ±8.0 | 48.6 ±2.2 | 44.7 ±0.9 | 95.7 ±1.0 | 353.3 ±12.3 | 0.834 ±0.142 |

**SI 6: The force-sensing cantilever spring constant and validation.** Force measurements were calculated from the visible deflection of the sensing cantilever and its spring constant. (a) The spring constant of the cantilever under a tip load was theoretically calculated using the moment of inertia of an ellipse (equation 1) and Euler-Bernoulli beam theory (equation 2) with constants from (b), and validated with AFM (N=6). The length constants were measured from calibrated images (N>5). The elastic modulus of PDMS, E, was measured with AFM (N=6). For measurements of microtissue force, the spring constant was calculated according to equations 1 and 3 to model the load at half the tissue thickness, $t_{1/2}$, from the top of the cantilever. The prestress, storage stiffness and phase lag measured without a load (ie. the noise floor) and an elastic standard (a polymerized 70x15μm strip of PDMS) are shown in c-e, respectively. The noise floor was much smaller than microtissue force measurements and remained unaltered throughout the examined strain range (N=6, linear regression, $p>0.5$). Measurements of the elastic standard were also invariant on the tested strain range (N=5, linear regression, $p>0.5$), and as expected, had near zero internal friction. Error bars represent the standard deviation.

$$I = \frac{\pi a b^3}{4} \tag{1}$$

$$k = \frac{3EI}{L^3} \tag{2}$$

$$k = \frac{6EI}{(L-t_{1/2})^2(2L+t_{1/2})} \tag{3}$$

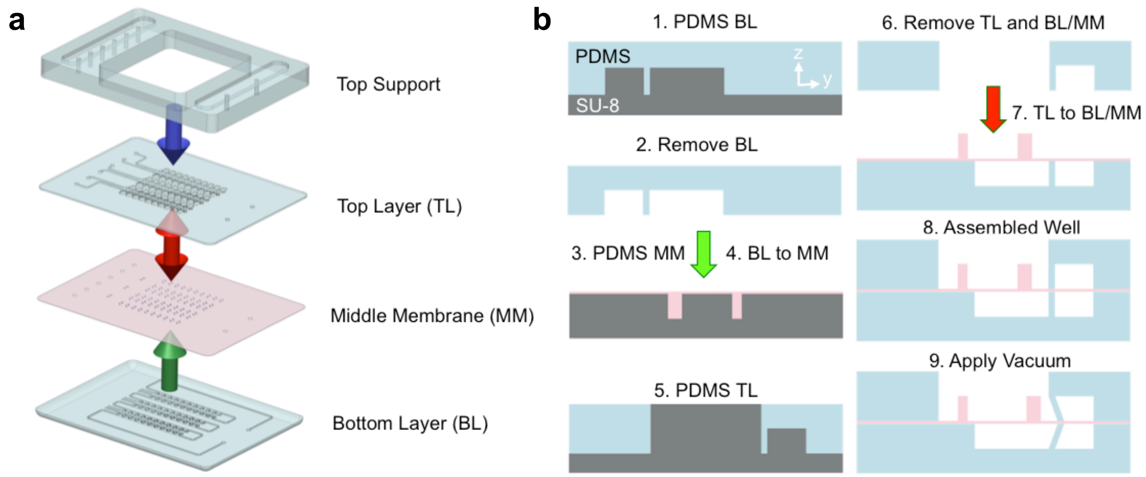


**SI 7: MVAS-force assembly.** The device consists of a top support and three photolithographic layers: 1) a top layer comprising the open-top microtissue wells and enclosed vacuum chambers; 2) a middle membrane with cantilevers; and 3) a bottom layer containing vacuum and empty bottom chambers. An exploded view is shown in (a) and cross-sections of fabrication steps of one well are illustrated in (b). For the bottom layer, PDMS is applied over top of the features on the SU-8 photolithography master and removed. For the middle membrane, PDMS, doped with Rhodamine B, is spin coated to a thickness of 30µm. Lastly, for the top layer, PDMS is spin coated to the top of the features on the SU-8 master. The bottom layer is then plasma bonded onto the middle membrane (green arrow) and the top support layer is bonded onto the top layer (blue arrow, not shown in (b)). The two are then removed from the middle membrane and top layer masters, respectively, and bonded together (red arrows). When a vacuum is applied, the middle membrane is deformed moving the cantilevers apart stretching the microtissue in plane.

**Supplementary Movie 1:** The MVAS-force device enables high throughput mechanical stimulation and force measurements of microtissues. In this movie, two microtissue cultures are simultaneously stretched in the MVAS-force device with a 0.25Hz sinusoidal vacuum. When a vacuum is applied, the cantilevers (on the left side) closest to the vacuum chamber move to stretch the microtissues while changes in microtissue tension are simultaneously measured by tracking the passive deflection in the opposing cantilevers (on the right side).
